# Conformational Space Profile Enhances Generic Molecular Representation Learning

**DOI:** 10.1101/2023.12.14.571629

**Authors:** Lin Wang, Shihang Wang, Hao Yang, Shiwei Li, Xinyu Wang, Yongqi Zhou, Siyuan Tian, Lu Liu, Fang Bai

## Abstract

The molecular representation model is a neural network that converts molecular representations (SMILES, Graph) into feature vectors, that carries the potential to be applied across a wide scope of drug discovery scenarios. However, current molecular representation models have been limited to 2D or static 3D structures, overlooking the dynamic nature of small molecules in solution and their ability to adopt flexible conformational changes crucial for drug-target interactions. To address this limitation, we propose a novel strategy that incorporates the conformational space profile into molecular representation learning. By capturing the intricate interplay between molecular structure and conformational space, our strategy enhances the representational capacity of our model named GeminiMol. Consequently, when pre-trained on a miniaturized molecular dataset, the GeminiMol model demonstrates a balanced and superior performance not only on traditional molecular property prediction tasks but also on zero-shot learning tasks, including virtual screening and target identification. By capturing the dynamic behavior of small molecules, our strategy paves the way for rapid exploration of chemical space, facilitating the transformation of drug design paradigms.

## Main

If we say that representation learning is the “sensory receptor” of artificial intelligence (AI) for complex entities, as the modulator for transforming human-readable information such as text, images, and sound into digital representations that can be understood by intelligent agents, then molecular representation learning is the interpreter to interpret the human-readable molecular representations (SMILES and Graph) into feature vectors for AI models to learn to perform drug discovery tasks. Molecular representation learning enables us to train models to understand molecular representations through pre-training tasks, thereby enhancing their performance in various downstream tasks. This approach is also known as meta-learning. Over the past few years, the scientific community has devised three prominent training strategies for molecular representation learning, i.e., self-supervised learning, supervised learning, and composite strategies.

Self-supervised learning is a popular training strategy that trains models to understand different formats of molecular representations (SMILES, IUPAC, InChI, Graph, 3D structure, molecular fingerprints, and even molecular image), such as SMILES-Bert^1^, ChemBERTa^2^, KPGT^3^, NYAN^4^, GraphMVP^5^, GEM^6^, Uni-Mol^7^, ImageMol^8^, etc. By leveraging contrastive learning or self-generated on diverse molecular representations, self-supervised pre-trained models can effectively extract features of molecular structures without requiring molecular structure labels. However, due to the requirement of learning a large number of unlabeled molecular structures during the pre-training stage, these molecular structures are likely to appear in the test set of downstream tasks as well. Hence, it becomes challenging to assess the generalization capabilities of self-supervised pre-training in the chemical space that is far from the training data. In terms of performance on downstream tasks, Sun et al. have pointed out that partial self-supervised pre-training strategies may not always exhibit significant advantages over non-pretraining methods^9^.

Supervised pre-training strategy is training with structure-properties projection or structure-caption semantic matching to capture molecular organic or pharmacological profiles, such as ChemBERTa-2^10^, MolT5^11^, CLOOME^12^, etc. Supervised pre-training strategies require different types of molecular attributes to form pre-training tasks, which can be certain classification or regression tasks, as well as retrieval or generation tasks of molecular descriptors or related biomedical data. The demand for labeled data leads to the major drawback of supervised pre-training strategies, which heavily rely on high-quality data. Besides, introducing molecular property information into the pre-training process raises concerns about the model’s generalization ability.

Considering the advantages and disadvantages of both self-supervised and supervised strategies, several composite strategies have been proposed to further enhance molecular representation learning, such as HelixADMET^13^. These strategies employ segmented pre-training stages, which involve additional supervised pre-training after self-supervised pre-training. At the self-supervised stage, they included tasks such as self-generation, molecular fingerprints generation, and 3D geometry prediction. At the supervised pre-training stage, they included tasks such as molecular properties prediction. This composite strategy enables such models to achieve superior performance in most downstream tasks.

To address concerns regarding potential chemical space biases and generalization capabilities in molecular representation learning, we propose a new hybrid contrastive learning framework. This framework combines inter-molecular contrastive learning with molecular similarity projection heads, enabling reliable molecular representation models to be trained on a miniaturized molecular dataset (39,290 molecules) without any experimental molecular properties. In this work, we trained a molecular presentation model using this strategy, named GeminiMol. This model incorporated both molecular geometric shape and pharmacophoric profiles of conformational space by learning the Conformational Space Similarity (CSS) of drug molecules. Following this, we examined our GeminiMol model’s performance in four major drug discovery tasks, i.e., virtual screening, target identification, Quantitative Structure-Activity Relationship (QSAR), and ADMET properties prediction.

### Descriptors of Conformational Space Similarity

The overall workflow of this study is illustrated in Extended Fig. 1. Firstly, we defined descriptors for conformational space similarity and established a contrastive learning framework based on molecular similarity. Subsequently, data collection was performed on a reduced set of molecules, and a model called GeminiMol was trained. The performance of the GeminiMol model was then evaluated on various molecular property prediction tasks, including independent datasets, out-of-distribution datasets, zero-shot drug discovery tasks (including virtual screening and target identification), real-world case studies, QSAR, and ADMET prediction, along with conducting ablation experiments. This workflow allowed us to progressively build and extensively test the GeminiMol model as a universal molecular representation model in various drug discovery scenarios, thereby expanding and deepening the research paradigm in molecular representation learning.

The similarity between molecular pairs provides the opportunity for contrastive learning, however, previous studies employed molecular pairs that were generated by modifying the molecular structure, such as MolCLR^14^ and iMolCLR^15^, rather than similarities between molecular pairs with pharmacological or physical significance. As illustrated in Fig. 1a, the 3D shape is a promising molecular similarity metrics that contains more pharmacological and physical information than the 2D structure and fingerprint similarity, therefore, 3D shape and pharmacological similarity is an optimal metric for contrastive learning. In GraphMVP^5^, GEM^6^, and Uni-Mol^7^, the authors primarily focused on 3D conformation as the pre-training task. In contrast, MolCLaSS^16^ incorporates static 3D shape similarity into the framework of contrastive learning. Static 3D conformation are employed in these approaches to facilitate molecular representation learning, however, the bioactive conformational space of drug molecules, plays an essential role in the drug-target binding^17,18^, the static minimal conformer used in previous works cannot represent the bioactive conformational space corresponding to the drug-target binding. In this study, we employ 3D shape similarity in the molecular conformation space as a metric for evaluating intermolecular similarity.

**Fig. 1.**
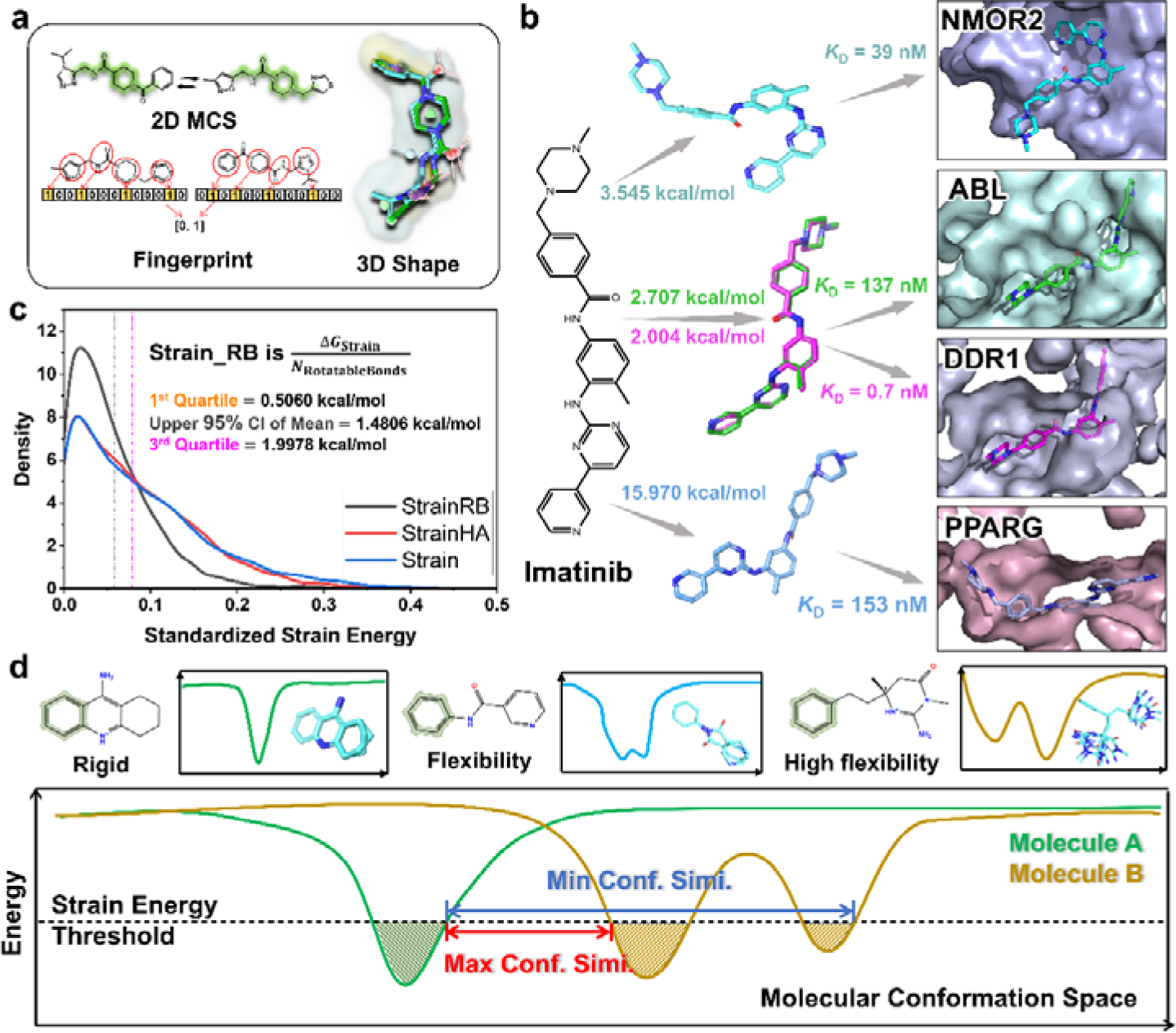
The definition of CSS descriptors of a pair of molecules. **a**, The different methods to present the molecular similarity. In this work, 3D shape means the molecular shape with pharmacophore features. **b,** The ratio between strain energy and the number of rotatable bonds exhibits the most optimal distribution curve. This density plot obtained by standardizing the ratio between strain energy and measurements of different molecular sizes, reveals the relationship between molecular size factors and strain energy. RB, rotatable bonds, HA, heavy atoms. **c,** Imatinib binds to four different targets with three different conformers. The energies in kcal/mol in this figure represented strain energy. The PDB IDs corresponding to the ligand structures from top to bottom are 3FW1, 3PYY, 4BKJ, and 6KTN. **d,** A schematic demonstrating the increasing complexity of a molecule’s conformational space with the number of rotatable bonds and the estimated approach of CSS.

Most small molecules contain multiple rotatable bonds, the rotations of which lead to conformational and energetic changes in the small molecules, resulting in a conformational space with several to hundreds of conformers. In particular, the small molecule is highly dynamic while binding into protein pocket, e.g., the most prestigious targeted drug, imatinib, binds to four different targets with three different conformers (as shown in Fig. 1b). Therefore, it is promising to incorporate conformational space profile of molecules into molecular representation model.

As illustrated in Fig. 1c, the strain energy distribution of 6,670 bioactive conformers of 4518 molecules in the protein data bank (PDB) ^19^ suggests that the number of rotatable bonds is a crucial factor in determining the molecular conformational space. With increasing numbers of rotatable bonds, the conformational space of a small molecule will exhibit an increasingly complicated distribution, containing more conformational clusters (as shown in Fig. 1d), which brings the potential to drug molecules for binding to multiple different drug targets. Designing a series of descriptors that can describe the CSS for contrastive learning is a reasonable strategy.

The key principle of the GeminiMol encoder is to extract the conformational space features based on the 2D molecular structures. To accomplish this objective, we oversee the model’s acquisition of conformation space features by utilizing various CSS descriptors. Towards this end, we devised a miniaturized molecular dataset (39,290 molecules) collected from six drug-like molecule databases, searched the conformational space of these molecules using two strain energy thresholds, and generated maximum and minimum similarities between pairs of molecules through pharmacophore and geometric shape alignment (the details were described in the *method* section). Additionally, we generated several CSS descriptors (as shown in Extended Fig. 2) by numerical transformation to combine the similarity scores obtained under the two different thresholds.

### Interpreting the contrastive learning framework of GeminiMol

In this study, we employed two different architectures to learn the CSS of molecular pairs, Cross-Encoder and Binary-Encoder (as illustrated in Fig. 2a). Cross-Encoder allowed the cross-attention weights between the reference molecule and the query molecule, this architecture was implemented by AutoGluon^20^ and ELECTRA^21^ model with chemical tokenizer. In the Binary-Encoder, the molecules are encoded independently using the same encoder (as illustrated in Extended Fig. 3) that was implemented as Weisfeiler-Lehman Network (WLN)^22^ and different projection heads. It is important to note that, both the query molecule and the reference molecule, as well as all similarity descriptors, share the same WLN network in the Binary-Encoder. The details of the tokenizer and Graph-Featurizers were described in the *method* section.

**Fig. 2.**
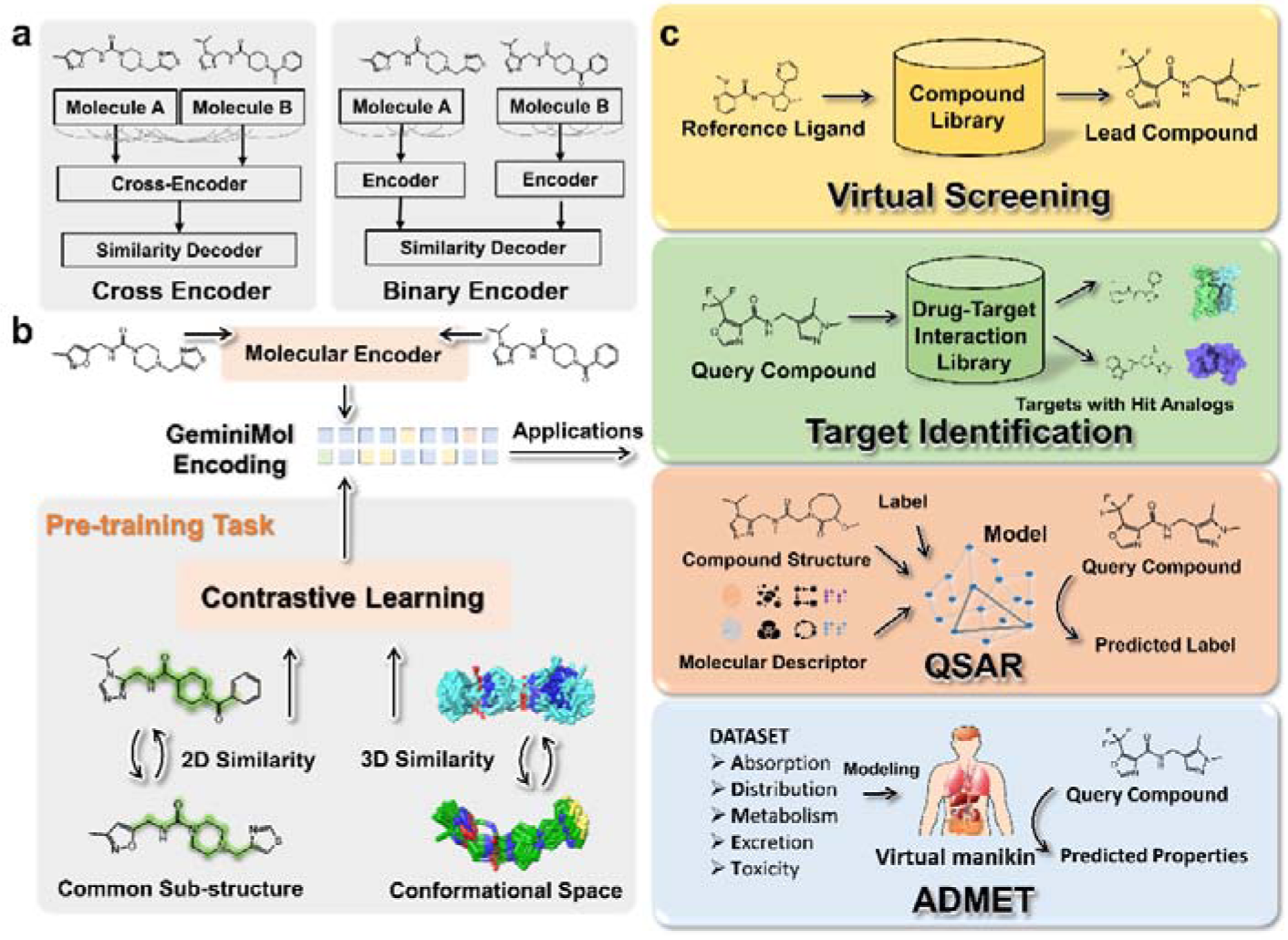
The training strategies of the molecular presentation models and their application tasks. **a**, Cross-Encoder and Binary-Encoder. **b,** The pre-training and application framework of GeminiMol. In the pre-training stage, both 2D and 3D molecular similarity were considered in our model, the 3D similarity was evaluated by PhaseShape, and the 2D similarity was calculated by the Maximum Common Substructure (MCS). **c**, our models were designed for four main drug discovery tasks, including virtual screening, target identification, QSAR, and ADMET prediction.

Based on these encoders, different projection heads were used for CSS descriptors and MCS similarities (as shown in Fig. 2b), the main purpose of these projection heads is to supervise the encoders in learning an informative molecular representation (named GeminiMol encoding). To evaluate the trained GeminiMol models, we applied the GeminiMol encodings to various downstream benchmark tasks as shown in Fig. 2c. We have developed benchmark datasets and corresponding baseline methods for various downstream tasks, including the DUD-E^23^ and LIT-PCBA^24^ for virtual screening, Target Identification Benchmark Dataset (TIBD) for target identification, target-based, cell-based, and ADMET-related compound properties data for QSAR and ADMET. See the *method* section for details.

The Cross-Encoder and GeminiMol (Binary-Encoder) were trained using a total of 8 million molecular similarity descriptors from 37,336 molecules in the training set, and then evaluated on both the cross and test sets. It is important to note that the cross set contains pairs of molecules that contain one molecule from the training set, whereas the test set does not contain any molecules from the training set. As shown in Fig. 3a, the comparison of the Spearman correlation coefficients of the models on the test and cross sets showed a strong correlation, indicating their robust generalization ability. To our knowledge, molecular pairs with significant differences in 2D structures and high similarity in 3D structures are rare but high-quality out-of-distribution samples. To investigate the impact of different components in the GeminiMol model on its learning ability, we performed ablation experiments on out-of-distribution data in the test set. As illustrated in Fig. 3b, reducing the number of layers and the dimensionality of node features, as well as removing rectifiers in the model, all have a detrimental effect on the model’s performance. Similarly, the same trend can be observed for CSS descriptors that undergo numerical transformations, as depicted in Fig. 3c.

**Fig. 3.**
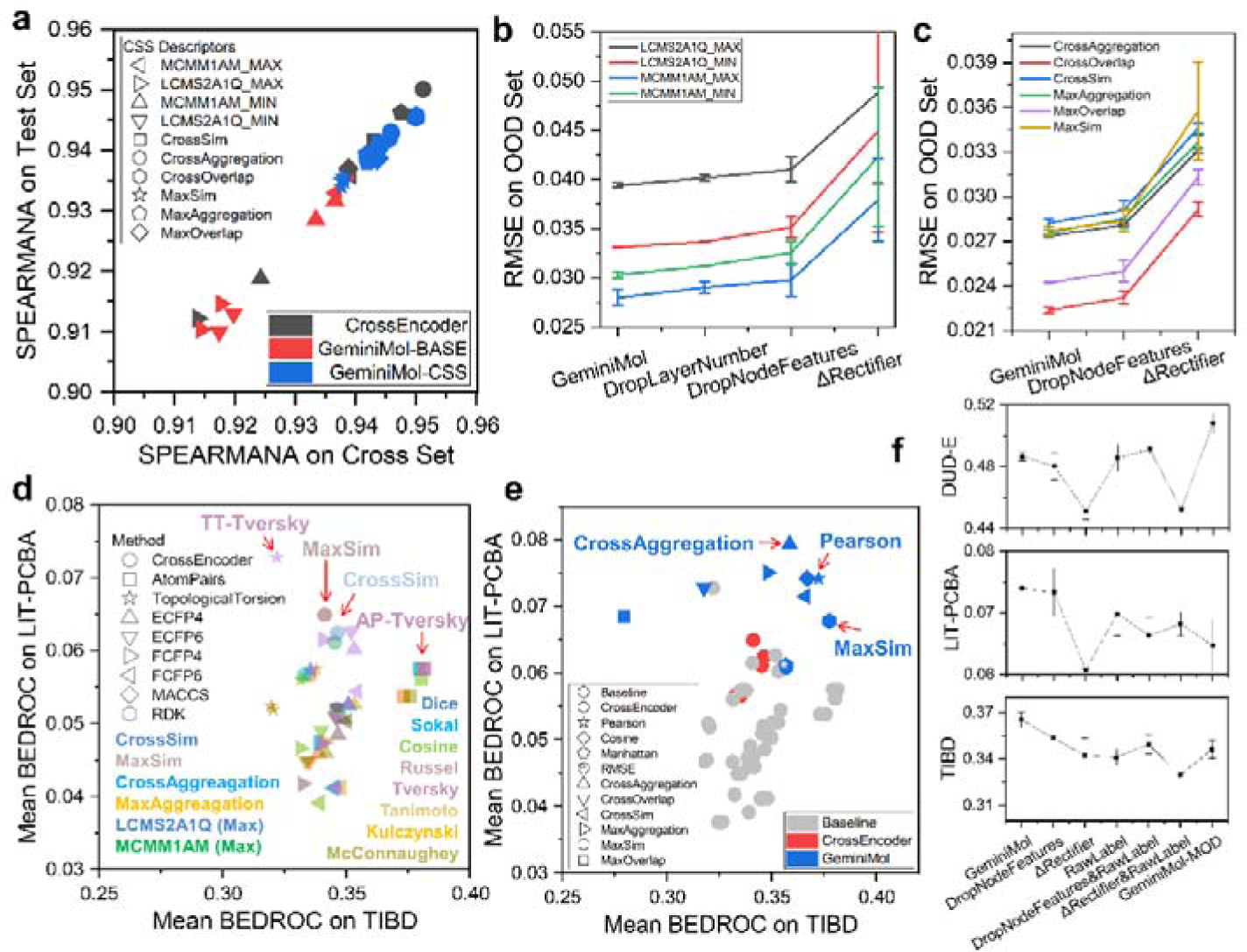
Learning the CSS descriptors and conformational space encoding using Cross-Encoder and Binary-Encoder. **a**, The correlation of performance between the cross and test sets reveals the high generalization capability of our models. **b,** Ablation results of raw CSS labels. **c**, Ablation results of transformed CSS descriptors. The GeminiMol model was subjected to four repeated training and testing experiments, while the other models underwent a minimum of two repeated experiments. The term DropLayerNumber refers to reducing the number of layers in the WLN network to 2. The term DropNodeFeatures indicates the reduction of node dimensionality in the WLN network to 1024. The term ΔReacifier denotes replacing the projection head to a simplification deep neural network. **d,** The performance comparison between Cross-Encoder with molecular fingerprints. The virtual screening performance is indicated by the BEDROC metric in the LIT-PCBA benchmark test, and the target identification performance is also BEDROC in the TIBD benchmark test. The choice of LIT-PCBA over DUD-E is primarily because the molecules in LIT-PCBA are derived from real high-throughput screening datasets. The model with the best balanced performance was pointed out with a red arrow. Molecular presentation methods and fingerprints were represented by various shapes, while similarity metric indices were depicted using different colors. **e,** The performance comparison of different molecular fingerprints, Cross-Encoder, and GeminiMol. **f**, Ablation results of GeminiMol models and CSS descriptors. The Pearson similarity metric was used in ablation experiments. The GeminiMol model was subjected to four repeated training and testing sessions, while the other models underwent a minimum of two repeated experimental and testing sessions.

### Towards Zero-Shot drug discovery using GeminiMol encoding

Zero-shot drug discovery, also known as virtual screening and target identification, involves identifying compounds from a compound database that exhibit intended activity and target specificity, relying solely on the structural features of one or a few molecules. This approach is particularly relevant in drug discovery, where active compounds are often limited in availability, necessitating the design of novel molecules with comparable activities using only a limited number of reference compounds. The molecular similarity searching serves as the primary methodology employed in zero-shot drug discovery.

In concept, molecules share similar conformational space also share similar biological activities, allowing us to predict the similarity of biological activities between molecules by comparing the similarity of GeminiMol encodings. Some projection heads used for training purposes can also be applied to compare the conformational space similarity of compounds, such as MaxSim, MaxOverlap, etc. Additionally, another approach is to directly compare the vector similarity between GeminiMol encodings, such as cosine similarity, Pearson correlation coefficient, Manhattan distance, and RMSE. It is worth noting that in this study, our miniaturized molecular dataset (39,290 molecules) exhibits few overlap with DUD-E^23^ (1,463,336) and LIT-PCBA^24^ (2,808,885). Therefore, the model has to learn the relationship between molecular 2D structures and their CSS while demonstrating excellent generalization ability to achieve better performance in downstream tasks.

We initially compared the performance of the Cross-Encoder and GeminiMol with PhaseShape on the virtual screening to investigate whether the incorporation of conformational space profile and contrastive learning improved the performance of PhaseShape, which was the shape alignment method used in the CSS data collection. As shown in Extended Fig. 4, many of the prediction heads, particularly the similarity between GeminiMol encodings, significantly enhanced the performance of PhaseShape on the DUD-E and LIT-PCBA benchmark datasets. Among the various encoding similarities, cosine similarity and Pearson correlation coefficient achieved the best overall performance.

To assess the differences between our models and prominent baseline methods, we evaluated the performance of 8 popular molecular fingerprints on the LIT-PCBA and the TIBD. We compared these results with those of the Cross-Encoder (as shown in Fig. 3d) and GeminiMol (as shown in Fig. 3e). The Cross-Encoder demonstrated comparable performance to molecular fingerprints, mainly attributed to the MaxSim and CrossSim predictors. However, its performance did not surpass that of molecular fingerprints. Surprisingly, the cosine similarity and Pearson correlation of the GeminiMol encoding exhibited balanced and superior performance across virtual screening and target identification compared to other methods. Further ablation experiments (as shown in Fig. 3f) demonstrate that removing the rectifier in the projection head leads to a significant detrimental impact on the model’s performance and robustness. To extract more information from the WLN network for molecular encoding, we have designed a variant of GeminiMol called GeminiMol-MOD, which enhances molecular encoding by reading the maximum and sum of each dimension of node features from the molecular graph. GeminiMol-MOD exhibits superior performance compared to GeminiMol on the DUD-E dataset. However, it shows some performance degradation in more realistic benchmark tests, such as LIT-PCBA and TIBD, as depicted in Fig. 3f. This could be attributed to the increase in feature dimensions, which leads to learning biases on molecular pairs with significant structural differences. This agrees with the observation that the DUD-E benchmark dataset has greater structural variations between positive and negative samples^25^.

### Applying GeminiMol encoding on scaffold hopping

To accurately represent the potential of molecules to exert their functions through similar targets and mechanisms of action, the molecular presentation model should effectively capture the underlying similarities between these molecules, even in cases where there are notable differences in their 2D structures. This aspect holds immense significance in the realm of practical applications, particularly in the pursuit of identifying novel active molecules with distinct scaffolds. In comparison to baseline methods, GeminiMol has demonstrated exceptional performance in zero-shot comparison of the molecular encoding, such as virtual screening and target identification (as illustrated in Fig. 3e). To gain insights into the underlying factors contributing to GeminiMol’s superior performance, we investigated its sensitivity to structural modifications and scaffold hopping of active compounds targeting three diverse targets.

We undertook a preliminary analysis to evaluate the impact of common ring-closure operations on the refinement of lead compounds, employing GeminiMol encoding, ECFP4^26^, and TopologicalTorsion^26^ fingerprints, as depicted in Fig. 4a. Our focus centered on two structurally analogous non-steroidal estrogen receptor modulators, specifically lasofoxifene and afimoxifene. Both compounds engage identically with the estrogen receptor, exhibiting comparable modes^27,28^ of binding (refer to Fig. 4a). Fingerprints suggest notable structural disparities, though such alterations remain within the predictive capacity of human analysis. In contrast, GeminiMol, acknowledging these compounds with nuanced structural changes as marked analogs, predicted a MaxSim value of 0.942, coupled with an encoding similarity surpassing 0.7.

**Fig. 4.**
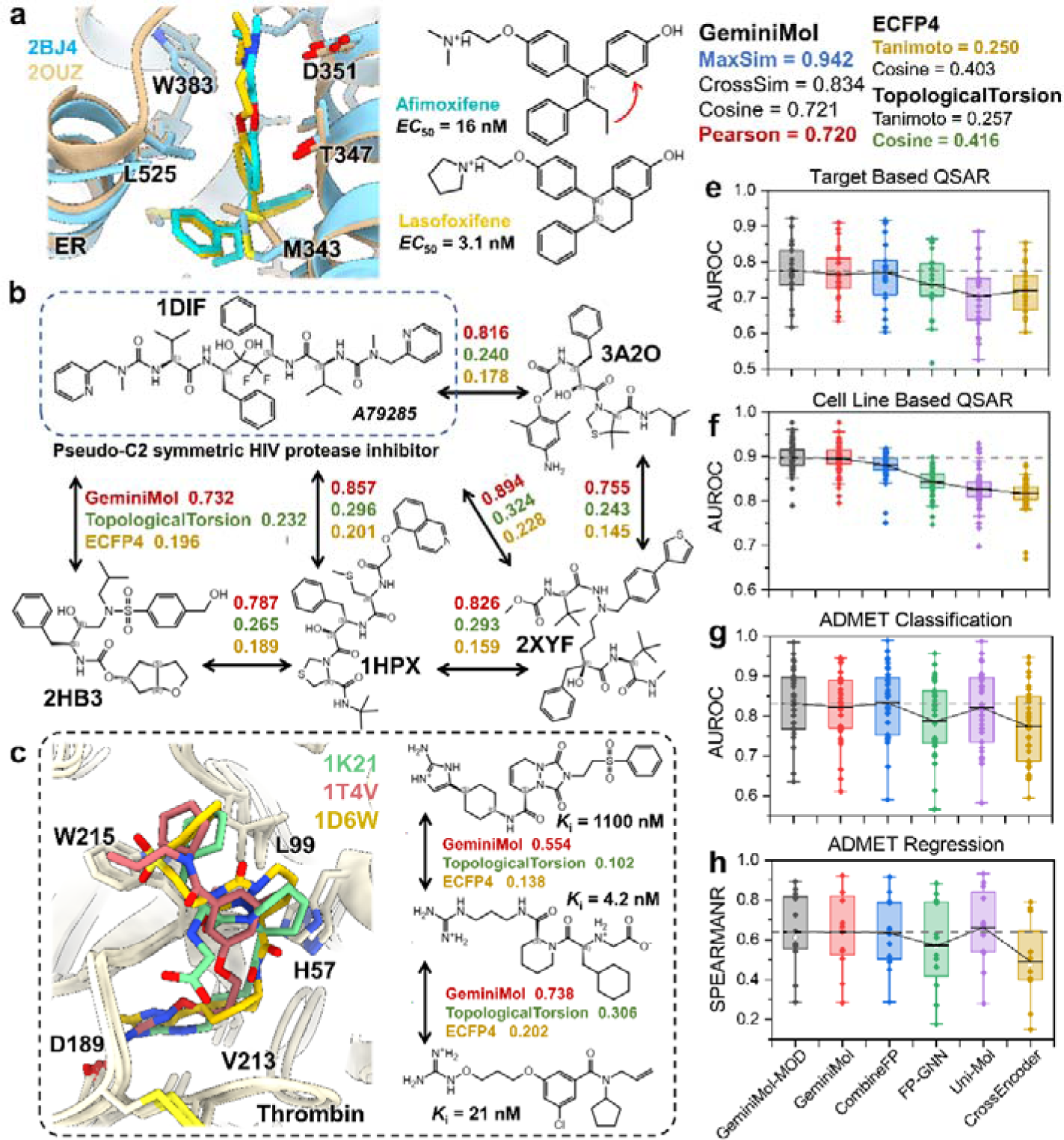
Interpreting and evaluating the GeminiMol encoding. **a**, Changes in molecular similarity induced by ring-closure. **b,** Intrinsic similarities of symmetric and non-symmetric HIV protease inhibitors. The Pearson similarity metric was used. **c,** Molecules with significant differences in their 2D structures can exhibit similar binding modes within protein pockets, while GeminiMol encoding also demonstrates similarities. The Pearson similarity metric was used. **e, f, g,** and **h** illustrate the comparison between GeminiMol and other baseline methods in four different downstream molecular property modeling tasks. The dashed line indicates the performance of the GeminiMol-MOD model.

The HIV protease has been a significant target in drug development with a comprehensive research history. Initially, research predominantly focused on pseudo-symmetric peptide inhibitors, evolving to diverse chemical scaffolds, while structurally distinct from the original peptide inhibitors, share a consistent mechanism of action. A comparative analysis was performed on the structural variety within these inhibitors, from the classical pseudo-symmetric inhibitor A79285^29^ against four non-symmetric inhibitors^30–32^. The molecular fingerprints suggested the 2D structural diversity of these molecules, however, it was observed that these molecules are in close proximity within the encoded space of GeminiMol (as shown in Fig. 4b). This observation highlights GeminiMol’s effectiveness in detecting intrinsic similarities within the conformational space of diverse molecules.

In a similar vein, we conducted a comparative analysis of the molecular structures of three thrombin inhibitors and their respective interaction patterns with thrombin^33–35^. This investigation reveals that despite substantial disparities in their 2D structures, the GeminiMol model is capable of identifying underlying similarities among these molecules. Notably, we observed a correlation between the activity of the inhibitors and their similarity to the most potent inhibitor, whereby lower similarity corresponds to diminished activity (as shown in Fig. 4c).

### Evaluating GeminiMol encoding on molecular property modeling

We compared the representation performance of GeminiMol and various molecular fingerprints on two different QSAR tasks, classification and regression ADMET tasks (see Extended Fig. 5). The results show that CombineFP (combined by ECFP4, FCFP6, AtomPairs, and TopologicalTorsion) performs the best among all fingerprint methods. In particular, QSAR tasks are divided into two categories: target-based and cell-based. The former is associated with specific targets, while the latter may involve multiple unknown targets and cytotoxicity mechanisms. In target-based QSAR tasks, GeminiMol-MOD, GeminiMol, and CombineFP demonstrate similar performance, with GeminiMol-MOD showing a slight advantage (as shown in Fig. 4e). In cell-based QSAR tasks, both GeminiMol models exhibit similar optimal performance, significantly outperforming CombineFP and other baseline methods (as shown in Fig. 4f). In the ADMET tasks, GeminiMol models perform comparably to CombineFP, achieving the best performance in classification tasks (as shown in Fig. 4g) but slightly underperforming the Uni-Mol model in regression tasks (as shown in Fig. 4g).

Overall, the GeminiMol model surpasses not only traditional molecular fingerprints but also FP-GNN^36^, designed specifically for QSAR tasks, and Uni-Mol^7^, which incorporates 19M molecular structures during pre-training, in terms of molecular presentation capability. This indicates that our proposed contrastive learning framework based on molecular conformational space similarity descriptors is an efficient strategy for training molecular representation models. Moreover, we investigated the performance of GeminiMol when solely utilizing the original GeminiMol encoding as input features for downstream tasks, while keeping the encoder fixed. Under this configuration of a fixed encoder, GeminiMol exhibits a slight decline in performance. However, it still outperforms the majority of baseline models in QSAR tasks and surpasses FP-GNN^36^ and certain molecular fingerprints in ADMET tasks (as shown in Extended Fig. 6). Remarkably, the fixed encoder was trained using only 37,336 molecular structures. This suggests that our training strategy endows the GeminiMol encoder with robust generalization capabilities. Additionally, incorporating a more diverse range of molecular structures in future research will further bolster the performance of GeminiMol.

### Related work

In recent years, somewhat inspiring works, such as GraphMVP^5^, GEM^6^, and Uni-Mol^7^, have utilized 3D structural information in molecular representation models. Besides, two similar conceptualized works, MolCLaSS^16^ and CLAPS^37^, incorporated physically or pharmacologically relevant molecular similarity labels in contrastive learning. To our knowledge, there is currently a lack of studies incorporating conformational space similarity into molecular representation learning.

## Discussion

The data quantity and quality pose significant challenges in AI for science. This study introduces an innovative approach to tackle this predicament, by harnessing highly precise techniques from molecular simulation to yield training data of superior quality for AI. The primary objective of this approach is to mitigate the scarcity of high-quality data in deep learning. By doing so, it facilitates the independent training of molecular representation models, while also deed as a supplementary data source for other strategies. Through our investigation, we have unearthed that incorporating descriptors of conformational space similarity can enhance the model’s representative capacity and efficacy in drug discovery. In contrast to similar works, we have taken great care in our framework design to avoid incorporating information that has already been considered in other representation methods, such as molecular fingerprints. Instead, we aim to explore novel means of representation, thereby paving the way for future model improvements.

Target-based drug design is the dominant paradigm in drug discovery in recent years. However, recent studies have shown that phenotype-based strategies have led to the discovery of the majority of drugs, and some target-based designed drugs also exert their effects through off-target interactions^38^. This finding suggests that there is still significant room for application in existing ligand-based drug design methods, including scaffold hopping and ligand similarity screening. Our work has revealed potential for improvement in the current molecular representation models, which inspires us to drive a paradigm shift in drug discovery. With the execution of initiatives such as Target 2035^39^, which focuses on the development of compound probes, similarity-based target identification is gaining prominence. Alongside advancements in molecular presentation techniques like GeminiMol, the bio-activities prediction and even the rational design of small molecules with specific target selectivity based on similarity networks between small molecules will become a new paradigm in future drug design. This study will also contribute to the fields of drug-target interaction prediction and drug side effect prediction.

Note that, the conformational space profile is not a panacea for drug discovery. For a portion of tasks, the 2D structure of a compound already contains sufficient information to establish structure-activity relationships, rendering the introduction of the conformational space profile inconsequential for these tasks. Additionally, the absorption, distribution, and metabolism of many drugs are closely related to the involvement of other biomacromolecules. Therefore, relying solely on information from small molecule conformational space is not an effective strategy for addressing such problems. GeminiMol aims to improve existing small molecule representation methods and provide a practical tool for other downstream tasks, such as protein-ligand affinity prediction and conditional molecule generation.

## Methods

### Strain energy analysis for bioactive ligands in PDB

The 6672 conformers obtained from the literature^40^ were first evaluated by energy calculations of the prime module of Schrödinger2021-1, and discarded the 2 conformers that were structurally implausible. Then, the strain energy analysis was performed by the strain energy rescoring program of Schrödinger2021-1. The solvent model used in the strain energy calculation procedure is the 4r distance-dependent dielectric (4RDDD) model and the global conformation searching method used is MCMM/Low-Mode in the MacroModel module of Schrödinger2021-1. The resulting strain energy values were divided by the number of rotatable bonds and the number of heavy atoms of the molecule and normalized, respectively, then, plot the density map in Fig. 1c. This was done to select an optimal factor to eliminate the effect of molecular size on the strain energy threshold.

### Schemes for CSS descriptors collection

In this study, a workflow was proposed to estimate the CSS descriptors for a pair of small molecules. As the schematic shown in Extended Fig. 7a, the CSS descriptors was calculated by three steps: 1) a pair of conformational ensembles were generated under the different strain energy thresholds and RMSD cutoffs. 2) superimposing the conformational ensemble of the query to reference molecule. 3) calculating their maximum and minimum molecular shape and pharmacological similarity to represent the similarity of the conformational space for a pair of small molecules. The strain energy thresholds for each molecule were determined by multiplying the number of rotatable bonds and the three threshold factors shown in Fig. 2c.

All the compounds in our dataset were sequentially performed the process of de-redundancy, ligand preparation, conformational searching and shape superimposition. The MarcoModel module of Schrödinger2021-2 was employed to search potential conformers of our molecule dataset. Both MCMM and Low-Mode/MCMM were used as search methods, and the RMSD used for eliminating redundant conformers was set to 1 or 2 Å. For each molecule, the energy window was multiplied by the product of strain energy factors and the number of rotatable bonds. The strain energy factors were three quartiles of the strain energy distribution of bioactive ligands of PDB (as shown in Fig. 2c), i.e. 0.5060, 1.4806, and 1.9978. The maximum number of steps was set to 2,000, and 100 per rotatable bond. The pharmacophore mode (-pharm) of PhaseShape^41^ was used for superimposing and similarity scoring the conformers of query molecule to the conformers of reference molecule. During the preliminary trials, we employed the CPU version of the PhaseShape module of Schrödinger2021-2 to superpose the generated conformers. During the training data collection phase, we used the GPU version of PhaseShape to speed up the computation.

### The construction of the miniaturized diversity molecular dataset

Considering the cost involved in sampling CSS descriptors as well as the diversity of the compound library, we constructed a miniaturized but high-quality compound dataset. As illustrated in Extended Fig. 7b, this dataset consists of six different subsets of compounds from the PDB, GPCR Ligand Libraries (GLL, a diverse library of molecules targeting the most predominant classes of drug targets), the active sets of Demanding Evaluation Kits for Objective *In silico* Screening (DEKOIS2), Enamine diversity libraries, Glide decoy molecules library, and the ChemBridge macrocyclic compounds library. The processed PDB ligands were downloaded from the literature^40^ (4461 unique ligands). The active ligands from the GPCR ligand library (16513 unique ligands), DEKOIS2.0 (2997 unique ligands), and diversity ligands from Enamine diversity collection DDS-10 (10214 unique ligands), Glide ligand decoys (1920 unique ligands), and ChemBridge macrocyclic compounds library (3156 unique ligands) were download from online databases. Then, we prepared them by the LigPrep module of Schrödinger2021-1. Molecular SMILES were generated by Schrödinger2021-1.

We further analyzed the structural diversity distribution of these compounds by Self-Organizing Map (SOM), as shown in Extended Fig. 8, which are fairly uniformly distributed in chemical space. This map was created using 96 most informative bits of the pairwise fingerprint of Schrödinger packages, the color gradient within cell population was set to 114 - 1,000, and the max population set to1,673. We demonstrated the relationship between the molecular structure and its SOM cell location in Extended Fig. 8 in a human-readable manner, where compounds between neighboring cells have a similar structure. In the preparation of SOM, the molecular fingerprints were computed by pairwise fingerprint and the 96 more informative bits were used for the inference of SOM.

### Collecting the CSS descriptors for training and evaluating GeminiMol models

Obviously, using CSS descriptors directly in virtual screening or target identification is an expensive approach, so it is difficult to compute CSS descriptors on the benchmark dataset to select the optimal strain energy threshold and RMSD cutoff for generating CSS descriptors. Therefore, a proof-of-concept study was performed to figure out the optimal strain energy threshold and RMSD cutoff for generating CSS descriptors. We randomly sampled 600,000 molecule pairs and calculated the shape similarity using the conformers sampled under the different strain energy and RMSD thresholds as described before. As shown in Extended Fig. 7c, the growth rate of the conformational space increases in a stepwise manner under different conformational searching principles, of which MCMM1AM (searched using MCMM under 1 Å and mean value of strain energy distribution) and LCMS2A1Q (searched using Low-Mode/MCMM under 2 Å and quartile of strain energy distribution) are two representative principles, and they belong to one of the two types of principles that produce the most and least conformations, respectively.

In addition, we analyzed the correlation between MCS similarity and the MaxSim, resulting in a vast majority of molecular pairs concentrated in an elliptical region as shown in Extended Fig. 9, whereas the upper right corner representing the region of high 2D similarity and high 3D similarity are apparent. We named these three areas as downtown, apparent and suburbia, respectively. Considering this imbalance distribution, we enhanced the sampling from suburbia area and reduce the sampling from downtown area in the subsequent data sampling.

We selected two different sampling principles (MCMM1A and LCMS2A) for CSS descriptors collection, and we mapped the CSS descriptors into a matrix (as shown in Extended Fig. 7d). As illustrated in Extended Fig. 7e, all the compound pairs were categorized according to two dimensions: 1) training (37336), validation (598), testing (1356), and crossover, and 2) downtown, suburbia, and apparent (as the ratio 0.30:0.65:0.05). The train, validation and test sets were divided by the molecular skeleton. For each pair, the indexes of the two molecules were exchanged to obtain the value of the symmetric position in the matrix, and abandoned this pair if the symmetric position is null. For the model using SMILES as input, i.e. Cross-Encoder, each SMILES in the dataset was randomized with a probability of 0.9 for data augmentation. We performed 10,000 rounds of molecular sampling on the training set, with 3,000 iterations per round, resulting in a dataset containing 183,391 positive samples with the MaxSim greater than 0.6. Considering the scarcity of samples with dissimilar 2D structures but similar 3D structures, as shown in the Extended Fig. 9, we extracted all samples from the training set with the MaxSim greater than 0.6 and not located in apparent regions, totaling 3,356,361 samples. The two sets of positive samples were then merged, and subsequently, negative samples were randomly selected from the sampled data with MaxSim less than or equal to 0.6 to complete the dataset, resulting in a total of 8,000,000 samples. A total of 4501 samples were extracted from the training set, where the MaxSim is greater than or equal to 0.75 and not located in apparent regions, to be used as an additional validation set. The validation set was sampled for two rounds, the test set was sampled for eight rounds, and the cross-validation set was sampled for twenty rounds. The molecule pairs excluding any molecules present in the training set that had a MaxSim greater than 0.6 and were not located in apparent regions were used as an additional independent test set.

### Tokenizers of the Cross-Encoder

The tokenizer used in this work is a Byte-Pair Encoding (BPE) tokenizer^42^, which vocabularies were obtained by segmenting the SMILES of the compounds in our dataset, and the last is a list of the common words that make up the SMILES, as determined by inspection by hand. These tokenizers all include at least the basic elements that make up a pharmaceutical small molecule, such as, C, N, O, S, P, c, n, o, s, F, Cl, Br, I, basic chemical bonds and isomer labels, such as, “=”, “@”, “/”, “\”, and “#”, as well as other special symbols, such as brackets and numbers.

### The details of the Cross-Encoder

We employed the AutoGluon^20^ framework to re-training the small version of the ELECTRA^21^ model (named SmELECTRA) using the hyperparameters as follow: learning rate is 0.001, weight decay is 0.001, learning rate schedule is cosine decay with 0.9, and the first 0.2 epoch was set to warmup steps. We confirmed ELECTRA’s superior performance in chemical languages in a previous study^43^. In these models, the CSS descriptor decoders are single linear layers. The training process was performed for a maximum of 20 epochs, and validated every 0.1 epoch, and up to three checkpoint models with the highest *R*^2^ on the validation set were stored. Finally, the top three models were fused by the greedy soup method and the final model was generated.

For CrossShape, we utilize the API provided by AutoGluon to extract conformational space encoding features. In AutoGluon, text is embedded separately and then concatenated together before being input into the model, allowing for cross-attention between texts. To prevent the model from simply counting the same tokens in both texts to predict similarity, we apply randomization to the input SMILES to enhance the data. This randomization is also used during the feature extraction process. Specifically, for the molecule from which we want to extract features, its canonical SMILES is placed in the position of the query molecule, while a randomized SMILES is generated and placed in the position of the reference molecule.

### Graph-Featurizers of the Binary-Encoder

We employed a specific graph featurization approach. The featurization process involves capturing diverse aspects of atoms and bonds within the molecular structure, which is crucial for accurately representing the chemical properties and interactions. For atom featurization, we utilized a base atom featurizers that incorporates multiple atom descriptors. These descriptors include atom type, atom hybridization, atom formal charge, atom chiral tag, whether the atom is in a ring, and whether the atom is aromatic. Each descriptor provides valuable information about the atom’s characteristics and its role in the molecular system. To combine these descriptors, we employed a concatenation featurizers, which concatenates the individual featurizers into a single feature vector. On the other hand, for bond featurization, we employed the canonical bond featurizers. This featurizers focuses on the bond type as the primary descriptor. The bond type indicates the nature of the chemical bond between two connected atoms and plays a crucial role in determining the molecular properties and interactions.

### The architecture details and training regimens for GeminiMol

The overall architecture of GeminiMol is illustrated in Extended Fig. 3. In the Binary-Encoder module, molecules are transformed into molecular graphs and subsequently encoded independently. The encoded vectors are then extracted using a specified readout function. In this study, we tested two main variations of the GeminiMol architecture, primarily differing in the way the encoded vectors are read out from the molecular graph and the structure of the molecular graph itself. For the GeminiMol model, the readout function is a multi-layer perceptron that takes node features as input and computes their mean value. For the GeminiMol-MOD model, we replaced the readout function with the weighted maximum and sum of node features. In the GeminiMol model, the WLN graph consists of 4 layers, and the node dimensionality is set to 2048. Due to CUDA memory limitations, in the GeminiMol-MOD model, we increased the number of layers to 12 while reducing the node dimensionality to 1024, as a replacement for the 2048-dimensional nodes in GeminiMol. In the ablation experiments, dropping the layer number means decreasing the WLN layers to 2, while dropping the node features means decreasing the WLN node features to 1024 dimensions.

The projection head of the Binary-Encoder employed in our study is designed to transform the learned GeminiMol encoding into meaningful output predictions. The network consists of a series of fully connected layers with non-linear activation functions and additional normalization and dropout layers to enhance its learning capabilities and prevent overfitting. The input to the CSS similarity projection head is two encoding vectors from query and reference molecules. The encoding vector first passes through a rectifier component, which expands the features to five times the dimension of the original encoding vector, batch normalization, and then reduces it to 1024 dimensions. Subsequently, the features gradually pass through linear layers of different sizes, reducing from 1024 dimensions to 128 dimensions and repeating this process three times. Finally, it outputs a value that is reset to the range of 0-1 through a sigmoid neuron in the projection head. The LeakyReLU activation function is added between all linear layers in the projection head. In the ablation experiment, a subset of models had the rectifier removed. For this subset, the encoding vector was directly reduced to 256 dimensions, and batch normalization was applied. Subsequently, the output was further reduced to 1 dimension after passing through LeakyReLU.

For all GeminiMol models, we use the AdamW optimizer and a learning rate of 0.0001 for Binary-Encoder training. The weighted MSELoss (Mean Squared Error Loss) function was used. In each training epoch, the training data was randomly rearranged and sampled 512 pairs per batch. We used two learning rate adjustment strategies during the training stage. First, the learning rate was gradually increased at the beginning of training. Second, we use cosine learning rate scheduling, which adjusts the learning rate according to the progress of the training steps to avoid falling into local optimal solutions. The starting learning rate of warmup is set to 0.1 times the original learning rate, which gradually increases to the original learning rate after 30,000 steps, and the minimum learning rate in the cosine learning rate schedule is 0.5 times the original learning rate, with a period of 10,000 steps. In addition, an early stopping strategy was used to determine whether the model converges by monitoring the performance on the validation set per 500 steps and terminating the training prematurely if there is no performance improvement in 50 validations iterations.

### The measurement of molecular similarity in this work

We introduce proprietary definitions of similarity when evaluating the similarity of three objects: the 2D structures and the 3D conformational spaces between molecules, and the coding vectors of the GeminiMol model, respectively. In this work, we denote the query molecule A and the reference molecule B using ***A*** and ***B***. We use ***N*** to denote the number of molecular bonds and the size of the encoded vector. In the following Equation 1 for representing the similarity of maximal common substructures (MCS), 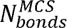 represents the number of bonds in the MCS, and thus, Similarity*_MCS,(AB)_* represents the proportion of the structure overlapping between the two molecules to the structure of the smaller molecule. Matching of element types is not strictly required when calculating the MCS.

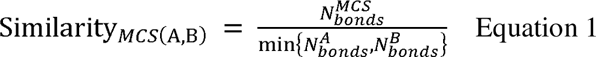

Besides that, the conformational space similarity for molecular pairs is represented by four metrics, which are represented based on the minimum and maximum similarity of the conformational space under two different strain energy thresholds. Among them, MaxSim is the maximum value of all similarities, MaxDistance is the minimum value of these similarities, MaxOverlap is one-half of the sum of MaxSim and MaxDistance, and MaxAggregation is one-half of the sum of the maximum similarities under both strain energy thresholds. In this paper, replacing “Max” with “Cross” in the names of these similarities means half of the sum of the similarities of molecules A to B and B to A. That is also referred to as the cross-similarity between the two molecules, which corresponds to the previous biased similarity. For cross-similarities, the similarity of A to B and B to A are identical, while for biased similarity, they are different. For example, Tanimoto similarity is a cross-similarity, while Tversky is a biased similarity.

Building upon the obtained conformational space encoding, the calculation of similarity between conformational space encodings using the vector similarity method described earlier can better describe the distances between molecules in latent space. This can be applied to various problems such as virtual screening, target identification, and scaffold hopping. For the encoding vector ***A*** of the query molecule and the encoding vector ***B*** of the reference molecule, we compute their RMSE similarity, cosine similarity, Pearson’s correlation coefficient, Euclidean similarity, Manhattan similarity, and Kullback-Leibler (KI) divergence similarity by using Equations 2-5 below.

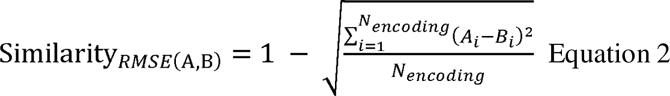

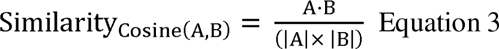

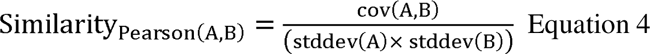

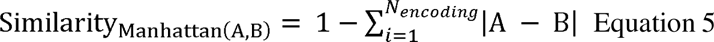

The Binary-Encoder independently encodes each molecule, allowing for a reliable calculation of the similarity between the encodings of two molecules, which in turn represents the similarity between the molecules themselves. This approach ensures a convincing representation of molecular conformational space encoding similarity.

### Collecting benchmark datasets for drug discovery

In this work, we selected the DUD-E^23^ and LIT-PCBA^24^ as the benchmark datasets of zero-shot virtual screening. Of these, DUD-E contained 102 targets and LIT-PCBA contained 15 targets. The molecular fingerprints were used in the decoy’s generation of DUD-E^23^ to avoid the production of “active” decoys, thus the superior performance of molecular fingerprints on DUD-E is well deserved, similarly, it same goes for the molecular docking and ligand similarity-based methods. In the LIT-PCBA^24^, both the active and decoy molecules were obtained from high-throughput screening, and the performance of all methods decreased substantially on this benchmark. For each target, DUD-E provides a reference molecule and LIT-PCBA provides several reference molecules. In the benchmark testing of LIT-PCBA, each target has multiple lead compounds that can serve as reference molecules. We calculate the similarity of all reference molecules to query molecules and select the highest similarity score as the final score for each query molecule.

In the scenario of evaluating virtual screening capabilities, DUD-E is a much easier task compared to LIT-PCBA, mainly due to the relatively fixed proportion of negative and positive compounds in DUD-E and the greater structural variations. However, it is very rare to encounter tasks as “standardized” as DUD-E in drug discovery, and it is common to encounter tasks as challenging as LIT-PCBA, in particular, compound activity cliffs, complex mechanisms of action, and large differences in the proportion of active compounds in different tasks. Therefore, LIT-PCBA is considered the main virtual screening benchmark test set in our study.

We processed the data in BindingDB to produce the zero-shot target identification benchmark dataset (as shown in Extended Fig. 10). In drug discovery, identifying the targets of active drugs is a challenging task, and the ligand similarity-based methods are the dominant approach in this field. However, the computational method is not necessary for molecules with apparently similar chemical structures (as illustrated in Extended Fig. 10a). As shown in Extended Fig. 10b, our benchmark dataset was produced by collecting decoy ligands with activity data entries over 300 and heavy atom numbers greater than or equal to 12 from BindingDB. For each ligand, we removed targets with binding affinities greater than 10 μM and also removed all ligands with MCS similarity greater than 0.5 to mimic real-world scenarios in which no similar structure compounds could be found from the bioactive database. Subsequently, we further refined the initial set of 176 decoy ligands by selecting only those ligands that targeted more than 70 receptors. This refined subset, consisting of 54 decoy ligands, was designated as the refined TIBD. The refined TIBD formulation comprises decoy molecules with diverse chemical structures, such as macrocyclic compounds like Ruboxistaurin, as well as Chlorpromazine (FDA-approved drug) with a single ring system, and natural product Quercetin. Using Dendritic molecular fingerprint and Tanimoto similarity, the heatmaps display that there is low similarity among most molecules in TIBD, indicating a high diversity of decoy molecules (as shown in Extended Fig. 10c). In our experiments, we used the decoy ligand to compare with all bioactive ligands in the decoy-specificity library and the targets extracted by matched ligands were used to analyze the ROC or enrichment factors (as shown in Extended Fig. 10d), so the final performance shows the performance of the different methods for discovering the targets but not bioactive ligands.

In addition, we have elaborated QSAR benchmark datasets corresponding to three different downstream tasks, including 21 single-target datasets from PubChem BioAssays^44^ database and LIT-PCBA^24^, 73 cell line datasets from the National Cancer Institute Developmental Therapeutics Program (NCI/DTP)^45^, and 34 classification and 12 regression datasets of the Adsorption, Distribution, Metabolism, Excretion and Toxicity (ADMET) properties from Therapeutics Data Commons (TDC)^46^ and a drug addictive dataset from the literature^47^. As mentioned above, datasets such as TDC^46^ and LIT-PCBA^24^ provide an avenue to evaluate molecular representation models in real-world application scenarios. Nevertheless, it is important to note that these datasets are predominantly influenced by target-based drug design approaches, resulting in a dearth of comprehensive mechanism-based phenotypic drug activity data. Most of the available QSAR datasets are derived from activity data specific to certain protein targets. Our benchmark datasets incorporated a total of 73 cell line datasets from NCI/DTP to investigate the performance of molecular representation models in multi-target QSAR tasks.

In this study, all training, validation, and test sets were divided using the molecular skeleton. For all datasets from TDC, we used the default scaffold split method to split the dataset (20% of data were held out for the test set). For other datasets from PubChem Bioassay, we used the Murcko Scaffold extractor provided by RDKit to identify the skeletons, the training and test sets were split according to the skeleton in a ratio of 7:3, where one-tenth of the data in the training set was used as the validation set. For the asymmetric validation embedding (AVE) division provided by LIT-PCBA, the training sets for per targets were also divided into training and validation using the molecular skeleton in a ratio of 7:3.

In this study, a variety of statistical metrics were employed to evaluate the performance of different methods. The area under the receiver operating characteristic curve (AUROC), the Boltzmann-enhanced discrimination of receiver operating characteristic (BEDROC), and enrichment factors with coefficients of 0.01%, 0.05%, 0.1%, 0.5%, and 1% were used to ranking problem, the AUROC and the area under Precision-Recall curve (AUPRC) were used to binary classification problem, the root-mean-square deviation (RMSE), the mean squared error (MSE), the mean absolute error (MAE), the Pearson correlation coefficient, and the Spearman correlation coefficient were employed to a regression problem. In most cases, BEDROC for ranking, AUROC for classification, and Spearman correlation coefficient for regression were the preferred metrics. Among them, BEDROC with α set to 160.9 was chosen, corresponding to EF_1%_ ^48^.

The zero-shot virtual screening and target identification were considered as ranking problem^48^. QSAR-based virtual screening tasks, i.e., seven QSAR datasets from LIT-PCBA, and the high-throughput assays data of CRF-R2, CBFn-RUNX1, NF-κ B, Peg3, ARNT-TAC3, A1 Apoptosis, uPA, Rango, Vif-APOBEC3G, RGS12, HSD17B4, and HADH2 from PubChem BioAssays^44^ database, NCI cell lines, drug addictive, and the classification tasks of ADMET were considered as the classification problem. The regression tasks of ADMET were considered as the regression problem.

### Collecting competitive baseline methods on these benchmark datasets

Over the past twenty years, various molecular representation methods have been developed to represent the features of pharmaceutical small molecules, including molecular fingerprints^49–51^, pharmacophores^52–54^, and molecular shapes^55–58^, which have continuously driven the technological development and the paradigm revolution of drug discovery. The molecular fingerprint is the most popular molecular representation method, which has been continuously updated since its proposal in the last century and is still the most dominant method of molecular representation^59^. Besides, the physics-based methods, including pharmacophores and molecular shape-based methods, such as AutoPH4^60^, ROCS^61^, and PhaseShape^41^, enable simultaneous presentation of both the geometric shape and pharmacophoric features of molecules, allowing for numerical calculations to compare the similarity between molecules. These similarity computation methods are highly interpretable, can be applied to a wide range of chemical structure types, and have enabled practical applications during the past twenty years. In this study, we introduce molecular fingerprints and PhaseShape as baseline methods to evaluate the performance of GeminiMol on zero-shot tasks.

To evaluate the virtual screening and target identification performance of our models, we first benchmarked the performance of 8 different molecular fingerprints (for all benchmark tasks) and 8 different similarity metrics (only for virtual screening and target identification) in RDKit^62^ on the benchmark datasets. This is done to preserve the bioactive conformers of the reference molecules. For Tversky similarity, we tried setting α to 0.01, 0.50, and 0.99, respectively. The performance of different combinations of molecular fingerprints and similarity metrics varies significantly across different targets and tasks. As researchers in drug discovery, we often desire a method that exhibits stable performance rather than having to speculate which similarity metric would work best for a particular project before it begins. Therefore, in this study, we aim to test specific combinations of individual molecular fingerprints and individual similarity evaluation methods, with the hope of obtaining a method that demonstrates stable performance across various tasks and can make accurate predictions. Besides, PhaseShape was also benchmarked on DUD-E^23^ and LIT-PCBA^24^, for the reference molecule structures required by PhaseShape, we utilize the protein preparation module in the Schrödinger software package for preprocessing.

The AutoGluon^63^, an AutoML framework, was employed for QSAR and ADMET benchmarks of different molecular representation approaches. We employ it for decoder network selection without stacking, bagging, and hyperparameters trials, with the purpose of comparing the performance of their optimal decoder to avoid the bias caused by the downstream decoder network. Candidate models include CatBoost^64^, LightGBM^42^, and neural network with PyTorch backend. The validation metric using AUROC for binary classification and RMSE for regression, respectively. It is worth noting that in this study, we did not use the hyperparameter searching of AutoGluon. Instead, we simply utilized AutoGluon to search for some commonly used model architectures without conducting hyperparameter tuning. The motivation behind this approach was to reduce the cost of benchmark testing while disregarding potential improvements from tricks.

In addition, we have incorporated a PropDecoder network act as decoder in QSAR and ADMET benchmarks, which has a similar architecture to the projection head in the GeminiMol network. The selection of validation metrics for PropDecoder depends on the type of dataset. For datasets with a positive/negative sample ratio greater than 3 or less than one-third, the AUPRC is used. Otherwise, the AUROC is employed. For regression tasks, the Spearman correlation coefficient is utilized. In terms of loss functions, for all classification tasks, the loss is calculated by taking the mean of BCELoss (Binary Cross-Entropy Loss) and MSELoss. Similar to the training process of the GeminiMol network, the AdamW optimizer is utilized during the training of PropDecoder. To adapt to varying dataset sizes, the complexity of the model, learning rate, and number of epochs are dynamically adjusted. For larger datasets, a larger learning rate, more complex network architecture, larger batch sizes, and fewer epochs are employed. Conversely, for smaller datasets, a smaller learning rate, simpler network architecture, smaller batch sizes, and more epochs are utilized.

Besides, FP-GNN^36^ and Uni-Mol were also benchmarked on all of the QSAR and ADMET tasks. For Uni-Mol, the data set splitting follows the same scheme as the molecular fingerprints and our models, the validation metrics and all hyperparameters are left at their default settings. For FP-GNN^36^, the data set splitting, validation metrics, and all hyperparameters are left at their default settings.

## Data availability

All related data are freely available from public sources. The molecular database, training data, and pre-trained models in this work were stored in Zenodo repository (10.5281/zenodo.10273480). The BioAssay ID and descriptions of the dataset can be found in Extended Table 1, while the statistical metrics for virtual screening and target identification benchmark tests are presented in Extended Table 2. Additionally, the statistical metrics for QSAR and ADMET benchmark tests can be found in Extended Table 3.

## Code availability

Source code for the GeminiMol model, trained weights, training and inference script are available under an open-source license at our GitHub repository (https://github.com/Wang-Lin-boop/GeminiMol), Neural networks were developed with PyTorch (pytorch.org), AutoGluon (auto.gluon.ai), and DGL-Life (lifesci.dgl.ai).

## Acknowledgments

We appreciate the technical support provided by the engineers of the high-performance computing cluster of ShanghaiTech University. Lin Wang also thanks Jianxin Duan, Gaokeng Xiao, Quanwei Yu, Zheyuan Shen, Shenghao Dong, Huiqiong Li, Zongquan Li, and Fenglei Li for providing technical support, inspiration and help for this work. We appreciate the developers of AutoGluon and Deep Graph Library (DGL). This work was supported by National Key R&D Program of China (Grant IDs: 2022YFC3400501 & 2022YFC3400500), Shanghai Science and Technology Development Funds (Grant IDs: 20QA1406400 and 22ZR1441400), Lingang Laboratory (Grant ID: LG202102-01–03), the National Natural Science Foundation of China (No 82003654), start-up package from ShanghaiTech University, and Shanghai Frontiers Science Center for Biomacromolecules and Precision Medicine at ShanghaiTech University.

## Contributions

W. and F. B. conceived the idea. L. W. designed the models, collected and tested all fingerprints, wrote the code, and performed the experiments. S. W. tested the baseline methods. S. W., H. Y., X. W., Y. Z., S. T., L. L., and S. L. contributed to data collection, case studies, and insightful discussion. All authors join in the preparation of the manuscript. The Large Language Models (GitHub Copilot, ChatGPT, and Claude) helped with a few touches to paper, and generated partial code.

## Supplementary information

**Supplementary Data**

**Extended Table 1. | General description of QSAR benchmark datasets used in this work.**

**Extended Table 2. | Statistical data of all molecular fingerprints, CrossEncoder and GeminiMol models in virtual screening and target identification benchmarks.**

**Extended Table 3. | Statistical data of all molecular fingerprints, CrossEncoder and GeminiMol models in QSAR and ADMET benchmark tasks.**

## Extended Data Figures

**Extended Figure 1.**
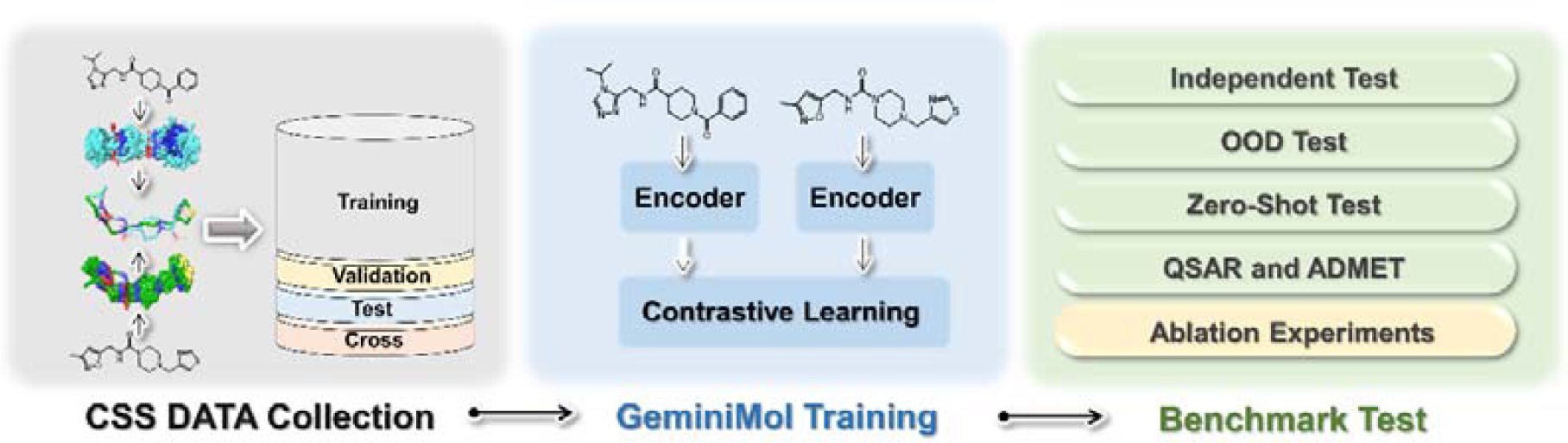
The overall workflow in this work. In this schematic diagram, the gray blocks represent physics-based data collection and proof-of-concept study, the blue blocks represent model training, and the green blocks represent benchmark testing on different downstream tasks.

**Extended Figure 2.**
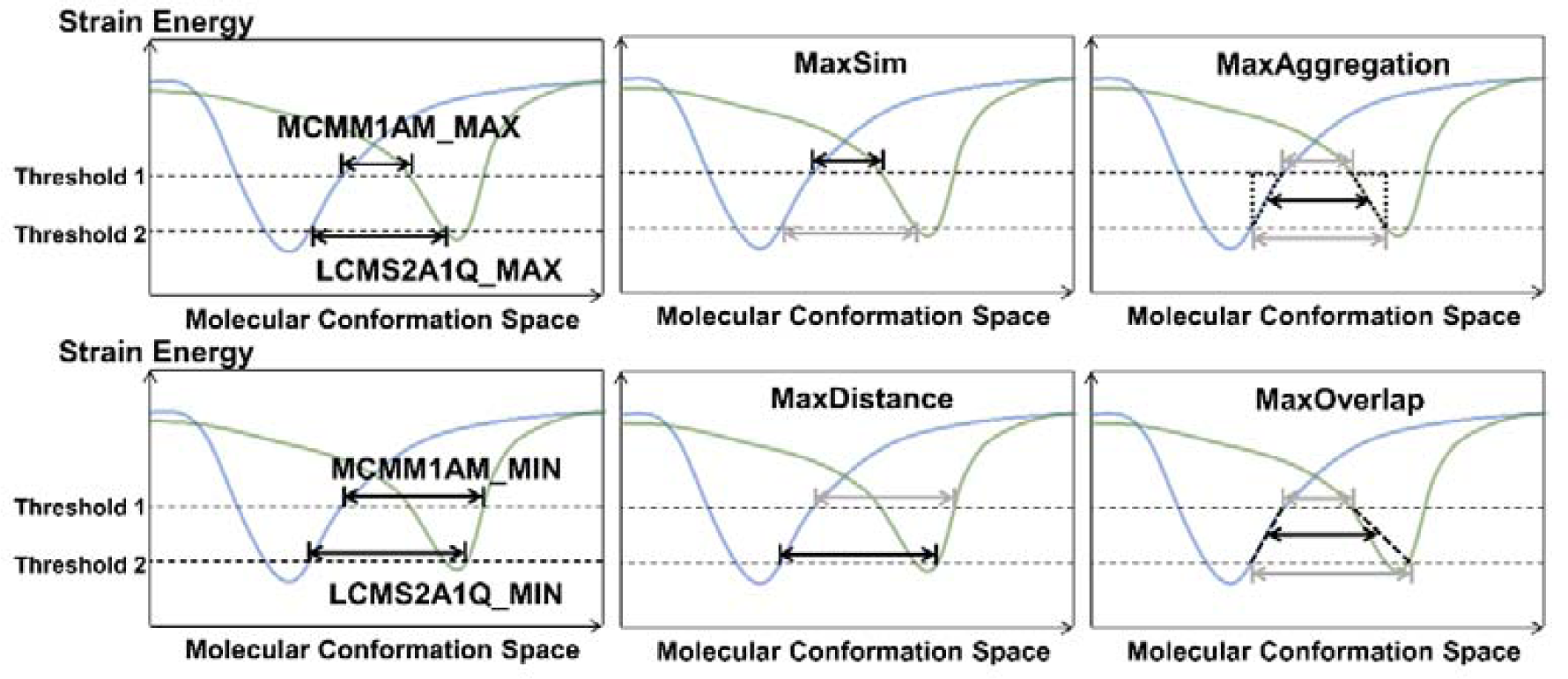
The CSS descriptors. The degree of similarity between the most similar conformations of molecules is represented by MaxSim, while MaxDistance represents the similarity of the most dissimilar conformations between molecules. The mean value of MaxSim and MaxDistance, referred to as MaxOverlap, is used to estimate the distance from any conformation of the molecule to the most similar conformation. The mean value of maxiaum similarity under two strain energies is regarded as the proximity of the conformational spaces of the two molecules.

**Extended Figure 3.**
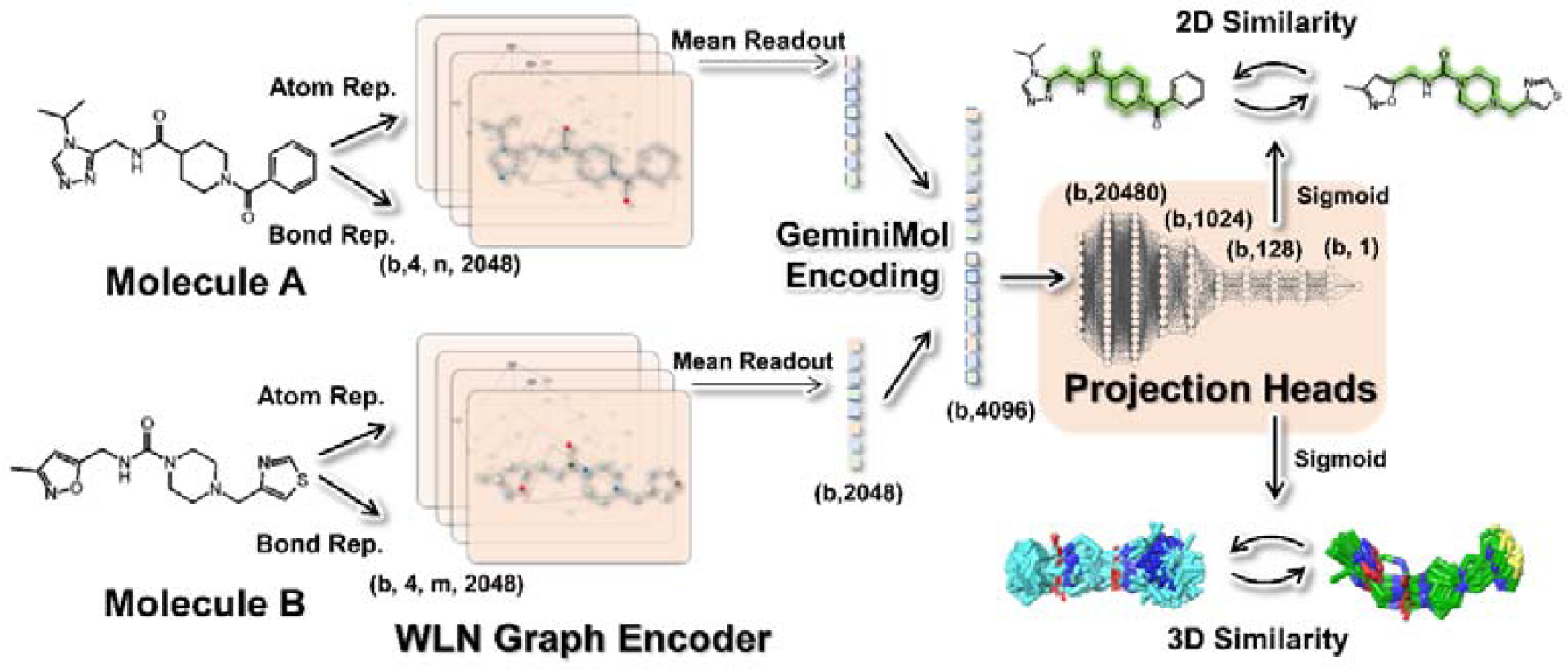
The Binary-Encoder architectures of GeminiMol. Molecules are fed into the network as graphs and encoded independently, where atoms are encoded as nodes and bonds are encoded as edges. Straight Arrows show the information flow among the various network components described in this paper. The highlighted 2D structures represent maximal topological common substructures (ignoring elemental differences). The dimensions of the feature vectors between the different network layers are labelled in parentheses.

**Extended Figure 4.**
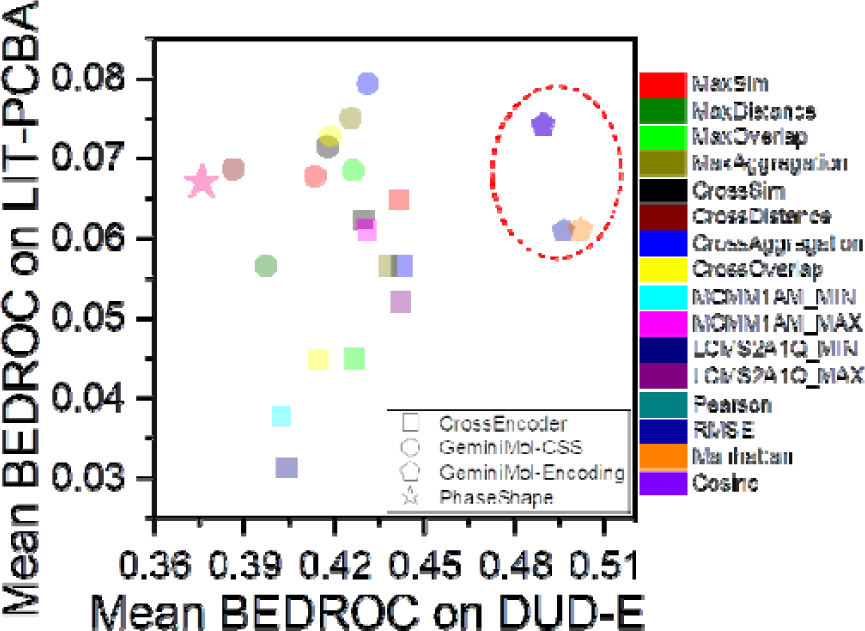
The virtual screening performance comparison of PhaseShape and our models. The portion highlighted in red dashed lines represents the similarity metrics of the GeminiMol encoding.

**Extended Figure 5.**
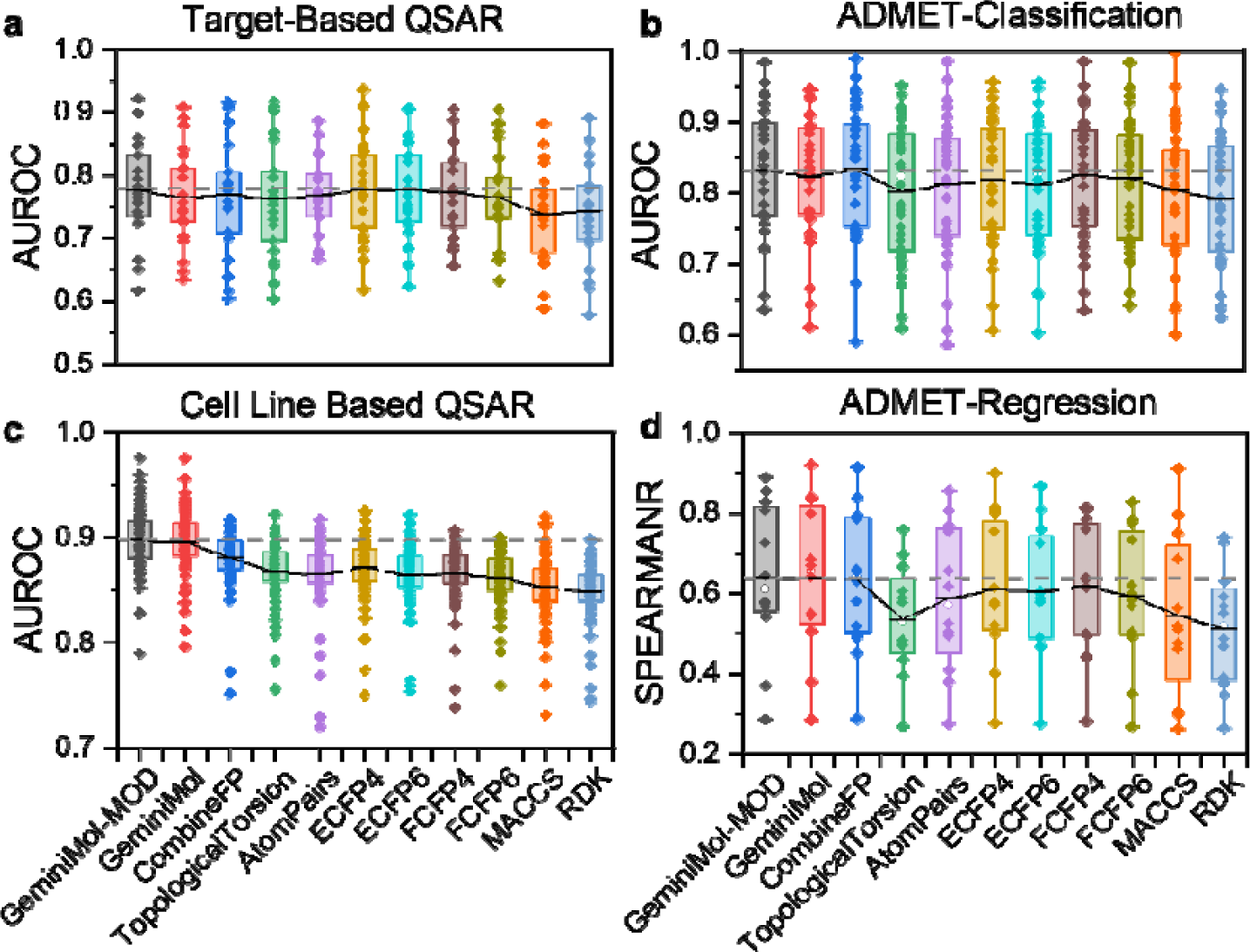
The comparative performance of the GeminiMol model and molecular fingerprints was evaluated across four different downstream tasks. **a**, Target-based QSAR. The data for this task was sourced from the PubChem BioAssay database and LIT-PCBA. All activity data exhibits dose-response relationships and is associated with specific target proteins. **b,** ADMET classification task. **c,** Cell-based QSAR task. This task focuses on QSAR modeling using data from 73 cancer cell lines obtained from NCI/DTP. **d,** ADMET regression task.

**Extended Figure 6.**
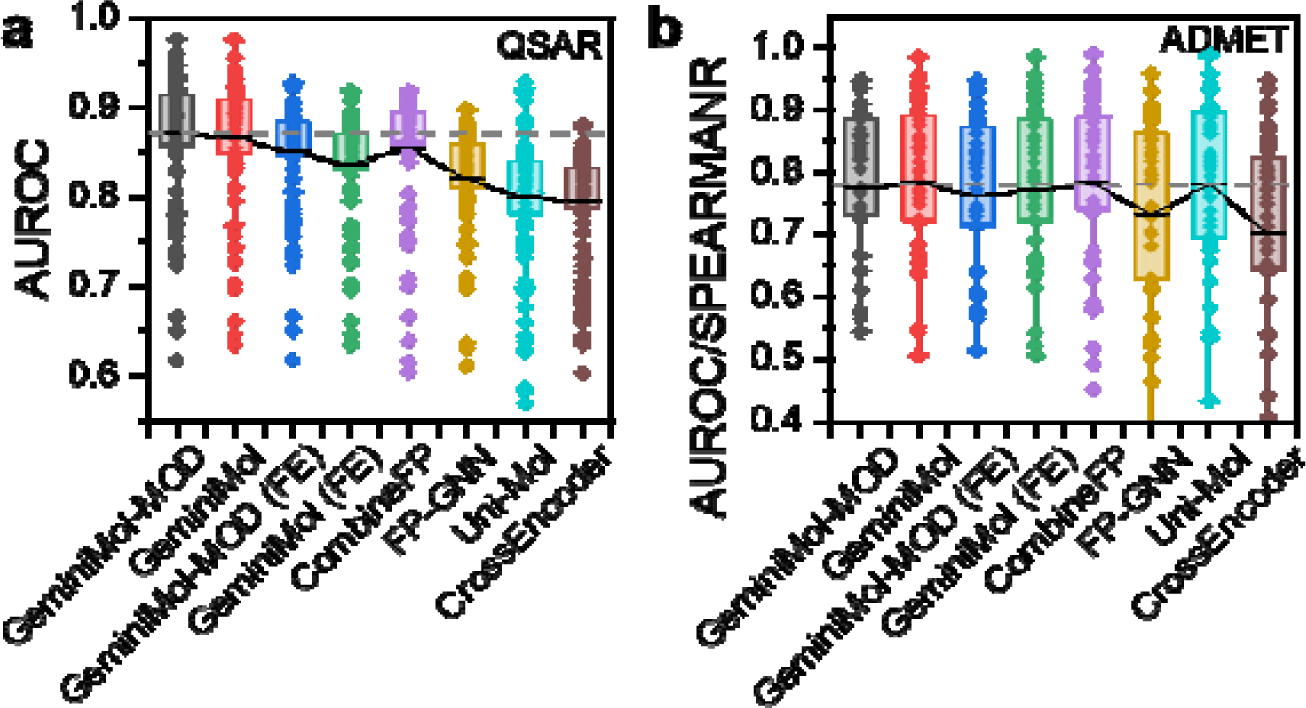
The impact of freezing encoder on the performance of GeminiMol for QSAR and ADMET tasks. **a**, The QSAR performance comparsion of freezing GeminiMol encoder with other methods. **b,** The ADMET performance comparsion of freezing GeminiMol encoder with other methods.

**Extended Figure 7.**
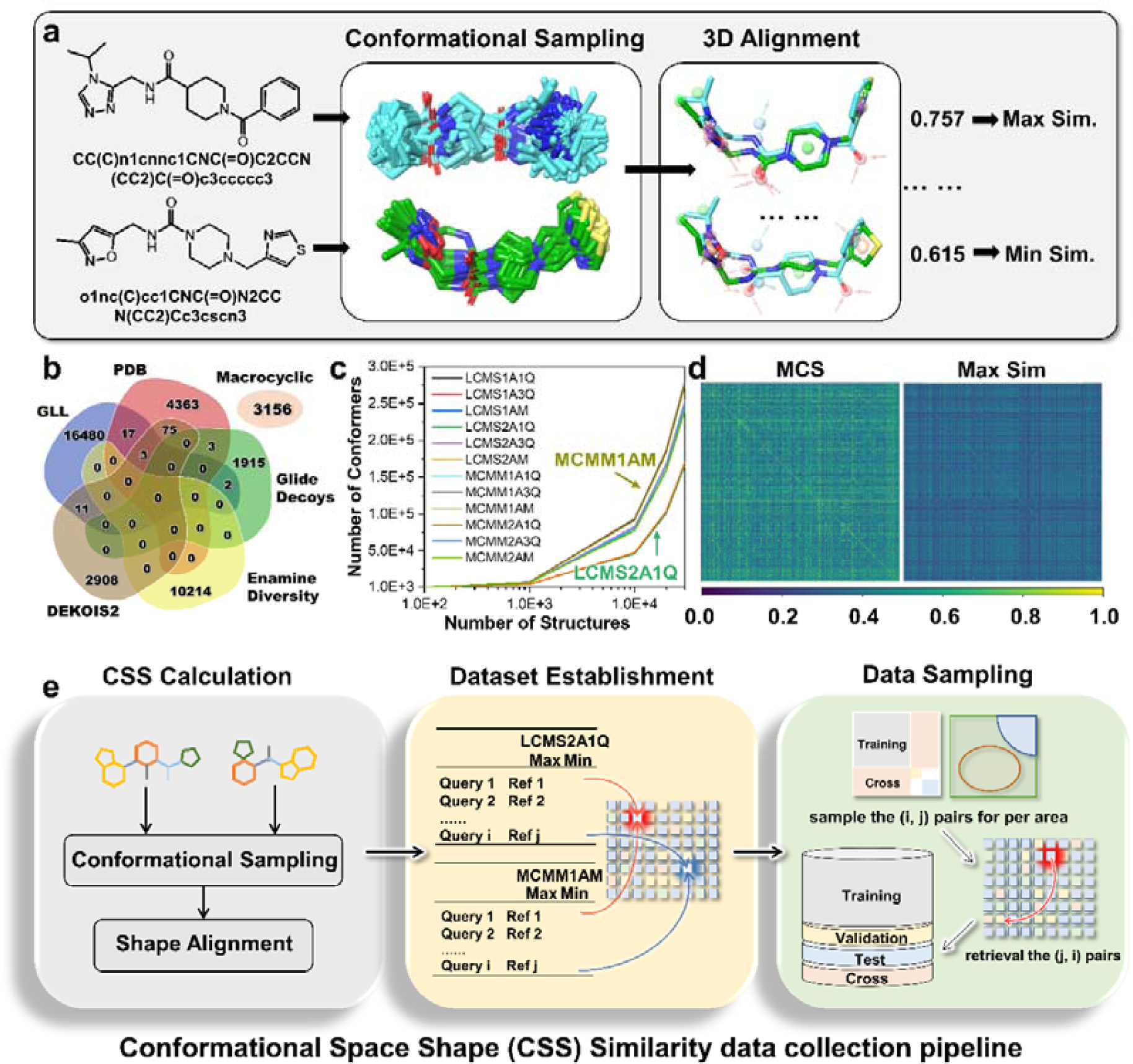
The data collection workflow of CSS descriptors. **a,** The flowchart illustrates the process of comparing the molecular CSS between two molecules. **b,** Source and size of molecular databases in this study. **c,** As the number of molecules increases, the differences in the number of molecular conformations under different sampling principles exhibit a stepwise growth pattern. The molecular order on the X-axis is fixed. **d,** Heatmap analysis reveals the diversity of the molecular dataset in this study. The two heatmaps on the left and right represent the MCS similarity and MaxSim similarity, respectively, which represent the 2D structural similarity and 3D structural similarity. **e,** Upon calculating the CSS, the obtained CSS descriptors under different conditions are then populated into a single matrix. Subsequently, symmetrically extracted CSS descriptors from the matrix are employed for the training, validation, and testing of the model.

**Extended Figure 8.**
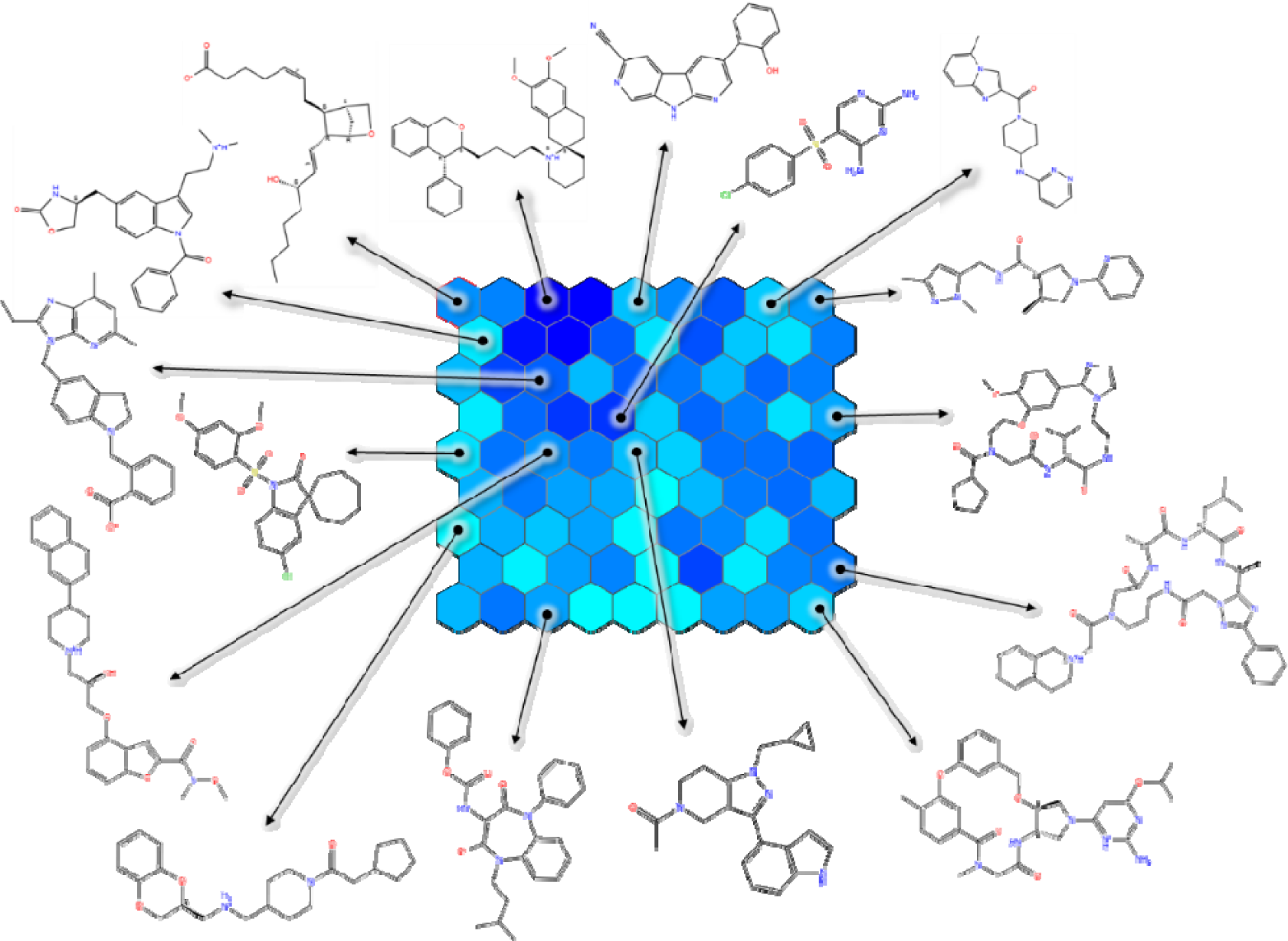
The SOM for visualizing the distribution of molecular structures in the compound dataset used in this work. The shown molecules were randomly selected from the cells. The SOM was created by 96 most informative bits of the pairwise fingerprint, and color gradient within cell population 114 - 1,000, the max population is 1,673.

**Extended Figure 9.**
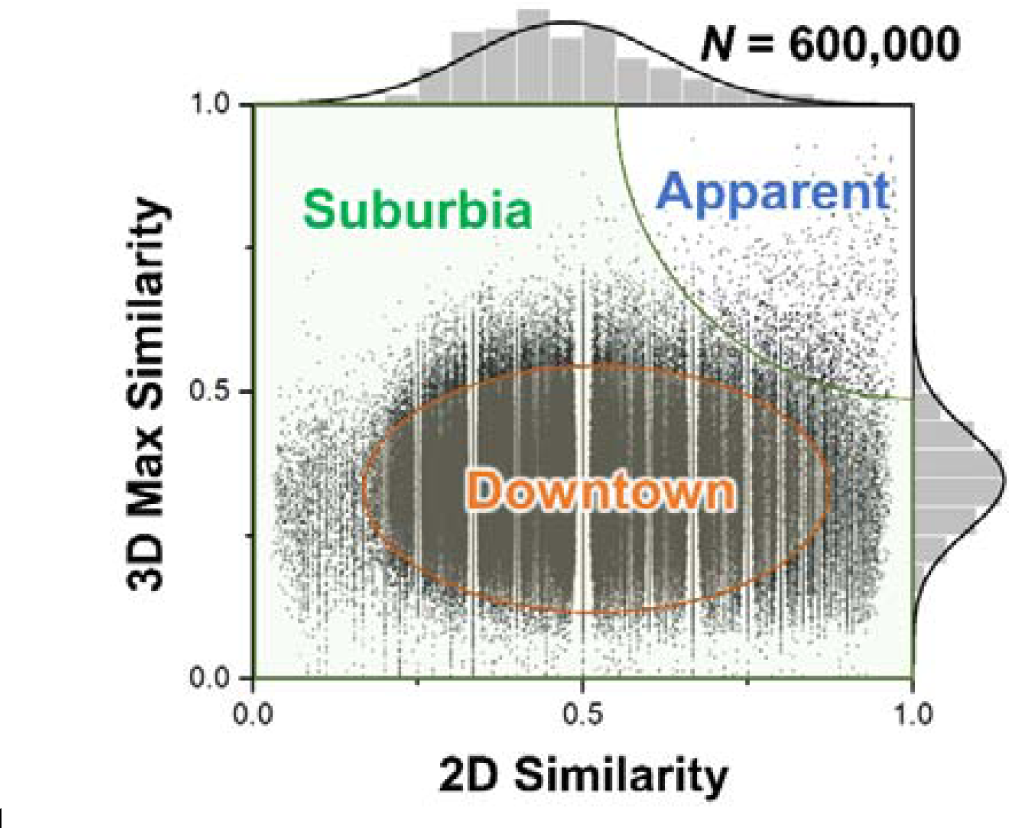
Relationship between MaxSim and MCS similarity scores for the molecule pairs in the preliminary experiment. The plot utilizes an ellipse to define a region with a high density of molecule pairs, and a sector to define a region with high MCS similarity and MaxSim values. During the training data sampling, different weights are assigned to these two regions, along with the remaining regions, to achieve balanced data sampling.

**Extended Figure 10.**
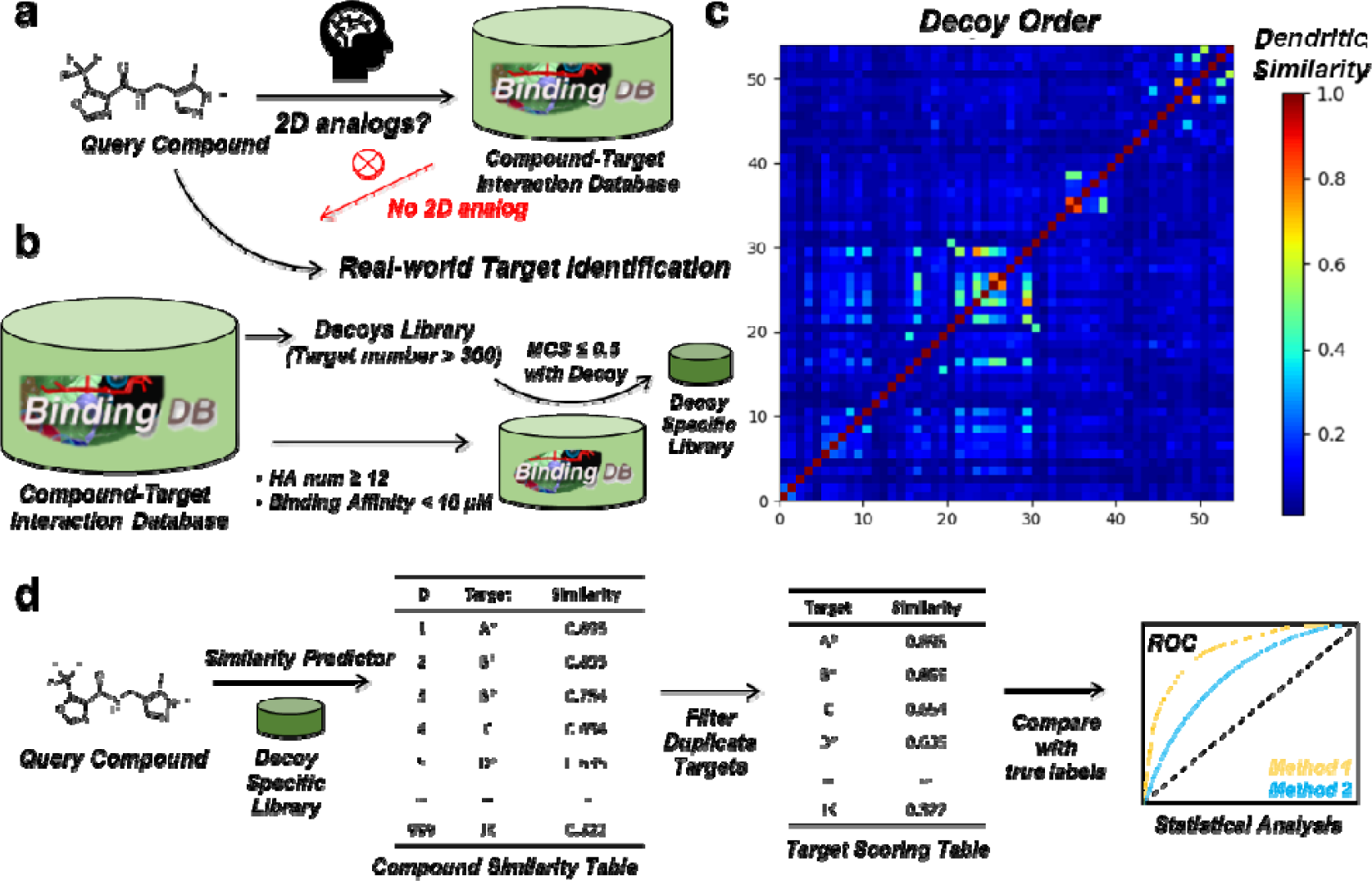
Motivation, process, and testing methods for the preparation of benchmark target identification dataset. **a**, For target identifying, query molecules with 2D analogues are easier to study. Therefore, ensuring that the compound-target relationship database does not contain 2D analogues of query molecules can better reflect the real-world scenarios in target identification tasks. **b,** Process for preparing the benchmark testing dataset. **c,** The fingerprint similarity heatmap of decoys. **d,** Proposed evaluation scheme for target fishing methods in this study. d, the heatmap of fingerprint similarities for all TIBD decoys. The similarity metrics is Tanimoto.

